# Switches in single-neuron excitability class can induce rich dynamical states in balanced networks

**DOI:** 10.1101/2025.02.28.640576

**Authors:** Moritz Drangmeister, Rainer Engelken, Jan-Hendrik Schleimer, Susanne Schreiber

## Abstract

Network states with rich dynamics and highly variable firing rates of individual neurons are prominent in experimental observations and thought to benefit complex information processing and learning. Such states have been proposed to arise from properties of network coupling, like a strong connectivity or slow synaptic dynamics. Here, we identify an alternative mechanism based on weak synaptic coupling and intrinsic cellular dynamics. We show that a switch in the cellular excitability class of action-potential generation (via a switch in the underlying mathematical bifurcation) results in super-Poissonian spiking variability in random balanced networks, where the recurrent network reinforces rather than averages out the single-cell effect. Information encoding is shifted to higher frequency bands and collective chaos in the network is enhanced when intrinsic cellular dynamics follow a saddle homoclinic orbit (HOM) bifurcation. Entropy production and attractor dimension peak close to the saddle-node-loop point that separates the two excitability classes, pointing to the qualitative nature of the transition as the origin of the effect. Because such switches are induced by common physiological variables, like temperature or extracellular potassium, and leave synaptic weights untouched, they offer a route to setting a network’s dynamical regime independently of plasticity. The robust effect links the biophysics of individual neurons to collective dynamics of large random networks, highlighting the relevance of single-cell dynamics for computation in physiological and artificial networks.

## INTRODUCTION

Cortical neurons fire more irregularly than models of balanced excitation and inhibition predict [1–7], and the discrepancy has consistently been attributed to properties of the synapses [8–12]. The irregular spiking activity emerging in models of balanced networks is associated with rich internal dynamics providing the substrate for complex adaptive computations [13, 14], and models that reproduce the higher variability observed experimentally accordingly rely on synaptic properties, including strong coupling, slow synaptic dynamics, or clustered connectivity.

Yet physiological properties of single neurons can profoundly affect the dynamics of their action-potential generation, shaping their spiking activity as well as the nature of interactions with other neurons [15]. In models of large networks, however, collective dynamics have been found to be remarkably robust to the details of the single-neuron description, and single-neuron properties are consequently often assigned a subordinate role [2, 16–18]. Here we demonstrate a mechanism that produces comparable high-variability states, yet originates in the properties of the single cells rather than in their connections. Specifically, we demonstrate that a qualitative switch in the single-neuron spike-onset bifurcation, i.e., the transition that underlies the initiation of regular spiking and is characteristic for a cell’s excitability class, can in balanced, random networks generate dynamics with strong spiking variability that exceeds that of Poisson firing. The switch occurs when the spike onset changes from the so-called saddle-node on invariant circle (SNIC) bifurcation to the saddle homoclinic orbit bifurcation (HOM), the latter of which has recently been identified as particularly interesting because of its effects on spike statistics [19, 20] and network synchronization [21–23].

The SNIC and HOM bifurcations are two out of three fundamental classes of neuronal excitability that determine how neurons capable of regular firing can initiate all-or-none action potentials [15]. They differ in the specific way, in which the resting state of a neuron loses stability when it receives a sufficiently strong input current to induce repeated spike oscillations. If the dynamics around this firing threshold are dictated by a SNIC bifurcation, firing rate increases gradually and uniquely as a function of the input current once the threshold is crossed (also termed class I firing, see [24] and [25]). A HOM neuron, by contrast, has two stable behaviours that coexist near the firing threshold, and noise can flip it between them – the coexistence that later gives rise to bursts. Dependent on the initial conditions at stimulus onset, neurons in response to a constant input current of given size either approach a subthreshold resting voltage (a fixed point) or fire regularly (on a limit cycle). Voltage can thus either linger near the stable resting state or transition to active limit cycle firing, before firing becomes the monostable state when the input current is increased further [26]. Accordingly, in the presence of noise or fluctuating input in the bistable range, burst-like firing patterns arise from stochastic switching between these two states, introducing longer pauses in firing between bursts (when dynamics are close to the resting state). Effectively, the bistability in the presence of fluctuations adds slower dynamics to the system [20].

Importantly, both bifurcations (SNIC and HOM) are topological neighbours in parameter space [21, 27, 28], separated by the codimension-two saddle-node-loop (SNL) point, so that transitions between these two excitability classes can be easily induced by small changes in biophysical neuron parameters. Theoretical studies, often supported by experimental evidence, have shown that changes in physiological parameters such as temperature elevations [22], increases in extracellular potassium concentration [29], alterations in ion channel expression [23] and even neuronal morphology [30] can induce HOM excitability in neurons, affecting not only their firing patterns but also the synchronization of mean-driven, pulse-coupled networks. While the latter finding can be understood from phase-oscillators theory, the effect of the HOM excitability class on large-scale, fluctuation-driven network dynamics remains mostly unexplored, in randomly balanced networks of the kind thought to be common in mammalian cortex [31–33].

The robustness of balanced networks to single-neuron detail has a clear basis. Most cellular parameters, like membrane time constants, thresholds, or gain, vary quantitatively. Such variation, including heterogeneity across a population, can often be absorbed into the self-consistent firing-rate distribution without qualitative change in the collective dynamics [34, 35]. A change in excitability class differs in kind: crossing the SNL point does not shift a parameter of the spike orbit, it alters its topology. This distinction, between quantitative variation and a qualitative change of the underlying dynamics, has received little attention in the fluctuation-driven balanced regime, precisely the setting in which single-cell properties are expected to be washed out, yet it offers a natural explanation for what we observe here. The intrinsic burstiness mediated by HOM excitability, and the resulting high spiking variability, do not merely survive embedding in a large balanced network. Rather than averaging the single-cell effect out, the recurrent network reinforces it, with significant influence on information filtering as well as on chaotic network states. Consistent with a topological rather than parametric origin, the effects are organized around the SNL point, with entropy production and attractor dimension peaking in its vicinity, and they persist both in conductance-based neuron models and for a Hopf spike onset. Cellular, physiological parameters thereby acquire direct relevance for setting a network’s state.

Because the excitability switch is driven by variables the brain routinely modulates, and because it leaves synaptic weights untouched, it offers a route to tuning a network’s dynamical regime, including its proximity to the edge of chaos, without interfering with the synaptic parameters that carry learning. The same mechanism links physiological perturbations to pathological network states, such as the bursting associated with febrile seizures or with potassium accumulation during epileptic activity. The insights provided in this study hence bear implications for understanding how the brain processes information, challenging the notion that rich internal network dynamics and strong spiking variability are only shaped by synaptic coupling. That cellular dynamics can set a network’s operating regime without touching its weights highlights the complexity of the individual unit as an under-used design dimension in artificial networks and neuromorphic hardware, spiking or otherwise.

## RESULTS

To investigate the impact of two excitability classes (defined by the SNIC and HOM bifurcations at spike onset) on the dynamics of balanced neural networks, we choose to implement mathematical neuron models with SNIC and HOM spiking dynamics based on their normal form reductions. This approach ensures the general validity of the results beyond the idiosyncrasies of specific conductance-based model choices. For both bifurcations we use the quadratic integrate-and-fire (QIF) neuron. In this one-dimensional model, the excitability class is determined by the value of the reset voltage after each spike, which captures the topological differences of the different spike orbits: SNIC for a reset to negative values and HOM for positive resets. Of special interest is the codimension-two bifurcation point, i.e. the point in parameter space where SNIC dynamics turn into HOM dynamics (also called the saddle node loop, SNL). Notably, when unfolding the SNL point, its normal form is identical to the QIF system with a reset around zero [36, 37].

In the following, we first provide details on the model and its behavior in balanced networks for SNIC and HOM cellular dynamics. We then analyze information filtering and the appearance of chaos and robustness of the observed effects in mixed SNIC-HOM networks.

Given a positive net input current to a neuron *Ī*, the quadratic non-linearity in the QIF sub-threshold voltage dynamics

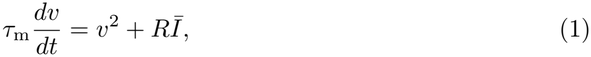

causes the membrane voltage to diverge to infinity in finite time, which represents the spike. In the canonical QIF model, equivalent to the theta neuron, the voltage is then reset to *v*_*r*_ =*−∞* after each spike. With *v*_r_ =*−∞* it corresponds to the center manifold of the SNIC bifurcation [38]. In addition to the input current, controlling the spike onset, the reset voltage can act as an additional bifurcation parameter which allows us to switch the onset-bifurcation to HOM spiking. Specifically, increasing the reset past the SNL point at *v*_r_ = 0 switches the neuron from SNIC to HOM excitability (Fig. 1a and b respectively).

**Figure 1.**
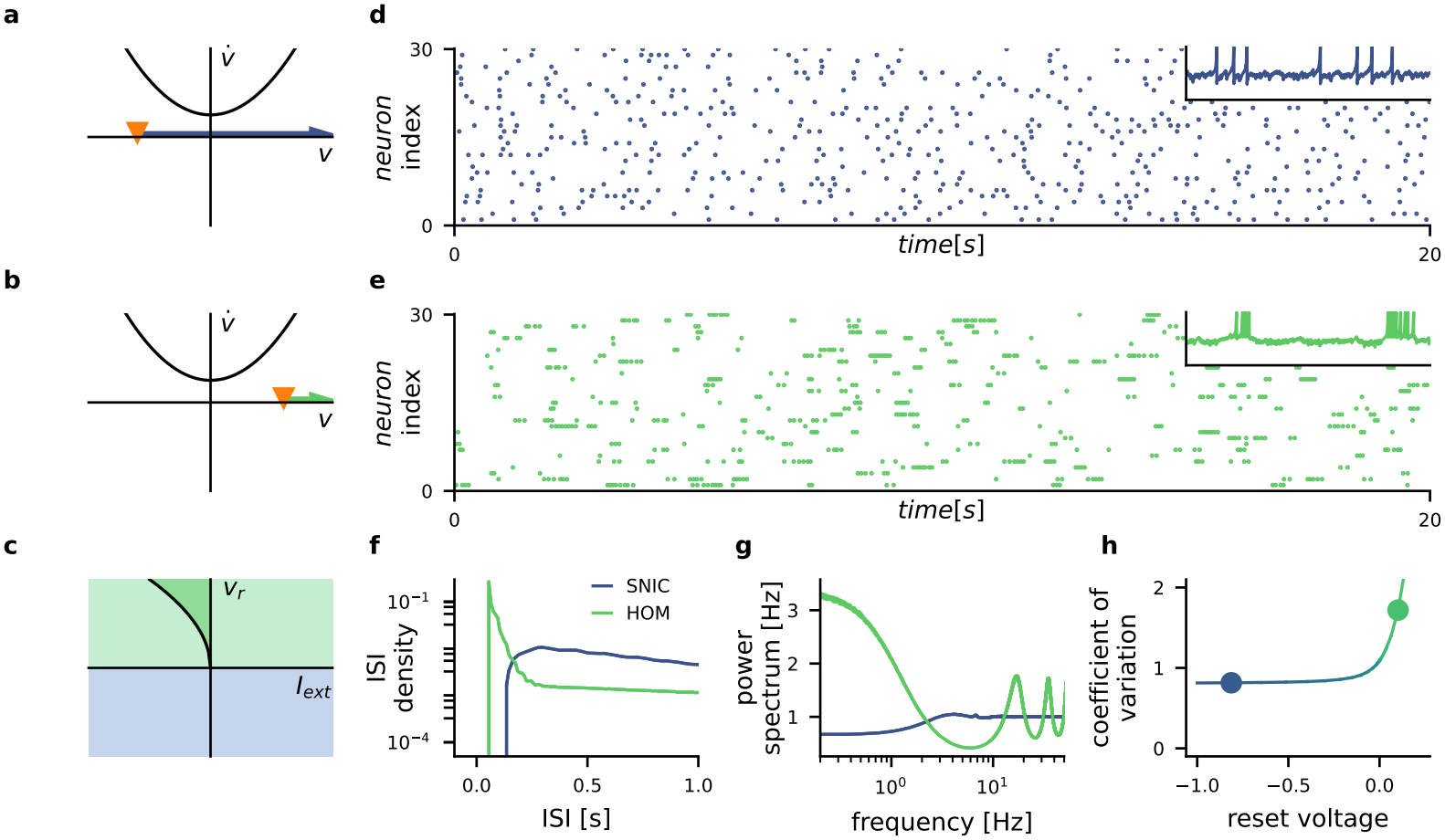
A switch in single cell excitability leads to burst-like activity in the network. Quadratic integrate-and-fire neurons transition from **a)** SNIC (blue) to **b)** HOM (green) spike onset when the reset voltage *v*_r_ (orange) is increased past the SNL point *v*_r_ = 0. **c)** The bistable region (shaded) of the HOM neuron is indicated in the full bifurcation diagram. **d)** Spike raster of a recurrent balanced network of SNIC neurons. **e)** Raster plot for identical network of HOM neurons. With the excitability, the network switches between different asynchronous states of spiking activity with Poissonian or burst-like dynamics. Inset shows example voltage trace. **f)** The ISI distribution of a neuron in the network reflects this switch, showing the HOM network with a peak at the fast intra-burst frequency and slower decaying tail. **g)** The spike train power spectrum of a neuron in each network. The slow switching of burst-like firing in the HOM network increases power at low frequencies. **h)** The coefficient of variation of the ISIs increases gradually with the reset, transitioning from sub-to super-Poissonian at the SNL point (*v*_r_ = 0)). The simulated networks consist of *N* = 1000 neurons, each with *K* = 50 inputs of strength *J*_0_ = 1. The reset is *v*_r_ = *−*0.8 for neurons in the SNIC network and *v*_r_ = 0.115 for HOM. The network activity is tuned to *ν* = 1 Hz.

The fundamental balancing mechanism of excitation and inhibition in the brain can be captured by a purely inhibitory network with constant excitatory input, offering a reduced model with fewer parameters [18, 39]. The balanced network analyzed consists of *N* = 1000 pulse-coupled inhibitory QIF neurons, with a fixed in-degree of *K* = 50 randomly drawn connections. In addition to some positive external bias current *I*_ext_, each neuron receives recurrent inhibitory feedback from the network

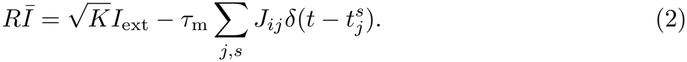

The resulting mean input into each neuron is slightly negative with spiking being driven by network fluctuations. In case of the HOM neuron, this implies the set point of the neurons to be in the bistable regime (Fig. 1c), where a stable fixed point and the spike limit cycle coexist. Note, that while fluctuations in the input can cross the spike onset of the neuron and cause it to fire, it can not alter the nature of the spike onset and its corresponding excitability class. We induce the switch from SNIC to HOM excitability by simultaneously changing the reset of all neurons. Because HOM neurons exhibit significantly higher firing rates, the external input current is used to tune network activity to a firing rate of *ν* = 1 Hz to rule out rate dependent effects.

In the following, we compare the activity of networks with identical topology and synaptic strengths which differ solely in the excitability class of the neurons involved (either all SNIC or all HOM) and *I*_ext_, as described above. Spiking activity in the HOM network is significantly more irregular (Fig. 1d, e). While cells in the SNIC network spike in the usual asynchronous irregular manner, characterized by a standard Poisson-like distribution of waiting times, cells in the HOM network typically exhibit long periods of silence interrupted by bursts of multiple spikes. Because of the shorter limit cycle period of HOM neurons, the peak in the inter-spike interval (ISI) distribution occurs at smaller intervals in neurons of the HOM network than those of the SNIC network (Fig. 1f). Also the spike train auto-power spectra differ significantly between the two networks. While the average spike power spectrum of the SNIC network shows Poisson-like firing with only a mild peak at the average firing rate and with the decorrelation of the balanced network resulting in a dip at low frequencies, the HOM network shows more pronounced peaks and increased power at low frequencies. The peaks occur both at the higher intra-burst firing rate and its harmonics. The power in the low-frequency range shows a Lorentzian-like power spectrum that reflects the slow switching rate between the two stable attractors, rest and bursting (Fig. 1g, Green line). The rich dynamics of the HOM network manifest in larger coefficients of variation (CVs). Fig. 1h shows the CV (averaged across neurons) as a function of the QIF model reset value *v*_r_ (color-coded), reflecting networks that change from SNIC to HOM dynamics (the point of transition, SNL, located at *v*_r_ = 0). The CV significantly increases when single-neuron dynamics become homoclinic. We further corroborated these results by simulating networks of conductance-based HOM neurons, finding the effect to remain qualitatively the same (Suppl. Fig. S3). The SNL point is located at a membrane voltage of *−*63.5 mV, confirming that the HOM simulations operate on the homoclinic side of the transition. The central role of the HOM bistability suggests a similar effect might arise from a Hopf spike onset – another excitability class and the other canonical route to bistability. Comparing it tests whether bistability as such is sufficient, independent of the specific HOM orbit. We demonstrate, that similar burst-like dynamics can be achieved in principle, but that the robustness and effect size are significantly reduced (Suppl. Fig. S4,S5).

### Shift to higher frequency encoding

Single-neuron information transmission is affected by dynamical systems properties, for example the PRC of the neuron [40]. Particularly, neurons with homoclinic onsets show sensitivity to high frequencies in their transfer functions [41]. A remaining question is if this property survives once neurons are part of larger recurrent networks contrary to the idea that the statistics of balanced networks, in particular second order statistics like spiking irregularity, are largely independent of the single-neuron model [16, 17]. We study the encoding capabilities of the network by adding a Gaussian input signal *s*(*t*) to a single neuron in the network and recording this neuron’s spike train *x*(*t*) as the response. Notably, the signal modulates the mean and not the variance of the input the neurons receives, which is known to alter the linear response [42]. In essence, this measures how much the neuron’s output tells us about its input, resolved by frequency, so that we can see which frequency bands it encodes well. From input and response, we estimate the mutual information rate, which quantifies the amount of information that a neuronal system encodes about an input signal it receives [43]. While the mutual information rate offers a single number, we use the frequency-resolved mutual information rate density *i*_LB_(*f* ) obtained from a Gaussian channel approximation, the integral of which was shown to offer a tight lower bound on the mutual information rate [44]

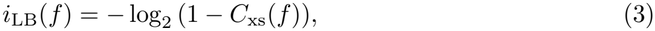

which allows an analysis of the filtering properties of the neuronal system. Here, *C*_xs_ is the squared coherence between signal and response of the neuron. To compare how the placement of a neuron into a recurrent network affects information encoding, this case is contrasted to renewal and Poisson approximations of the recurrent network input (see Methods). Overall, the information content per frequency bin of a single neuron in the network is low in all cases, as expected, but its distribution across frequencies differs depending on the excitability class. The spike response of single SNIC neurons has been reported to act as a low-pass filter, favoring the encoding in the neuron’s firing rate [45], which means it is capable of locking to low-frequency information in a stimulus, (Fig. 2g). This property of the gain is shared by HOM neurons, but their ability to lock to stimuli also extends to higher frequency ranges (Fig. 2g). In addition and in contrast to the SNIC neurons, the HOM neurons, due to their inherent burstiness, produce a larger amount of intrinsic noise in the low-frequency range (see Fig. 2h). The increased power in the intrinsic slow frequencies of the HOM neuron suppresses information encoding at the low end of the spectrum compared to the SNIC case (Fig. 2e). This effect is strengthened by the recurrent network, because it boosts the intrinsic burstiness (Fig. 2f). At higher frequencies around the limit cycle frequency, however, the encoding capabilities improve. While this effect is small for the Poisson-driven neuron, it effectively shifts the filtering behavior towards a band-pass filter. The recurrent network input reinforces this shift significantly, as it also reinforces the burst-like activity. The renewal approximation replaces the recurrent network with a single neuron driven by surrogate input whose statistics are iterated to self-consistency, so that the neuron’s own output statistics match those of its input (see Methods) [46]. The surrogate input carries the firing statistics of the network but none of the correlations that the recurrent connectivity imposes between neurons. It therefore contains no network interactions beyond this self-consistency – yet it reproduces the full network statistics quantitatively. This agreement is itself informative: it shows that the amplification of burstiness arises from the self-consistent single-neuron response to network-like input rather than from the correlations generated by the network structure, locating the origin of the effect at the single-neuron level. Further considering the additional effect of network parameters on the firing variability, we observe an even more drastic increase of the CV at the SNL point for weaker coupling (Fig. 2i). This observation stands in contrast to previous studies on comparable networks, where burst-like activity was induced through an increase in synaptic strength [9, 10]. The disparity between these results highlights the fundamental difference between a mechanism based on single-neuron excitability and a network coupling effect, providing an access point for experimental validation. In agreement with recent studies [47, 48], we also found that QIF networks exhibit population oscillations for high in-degree (see Suppl. Fig. S6).

**Figure 2.**
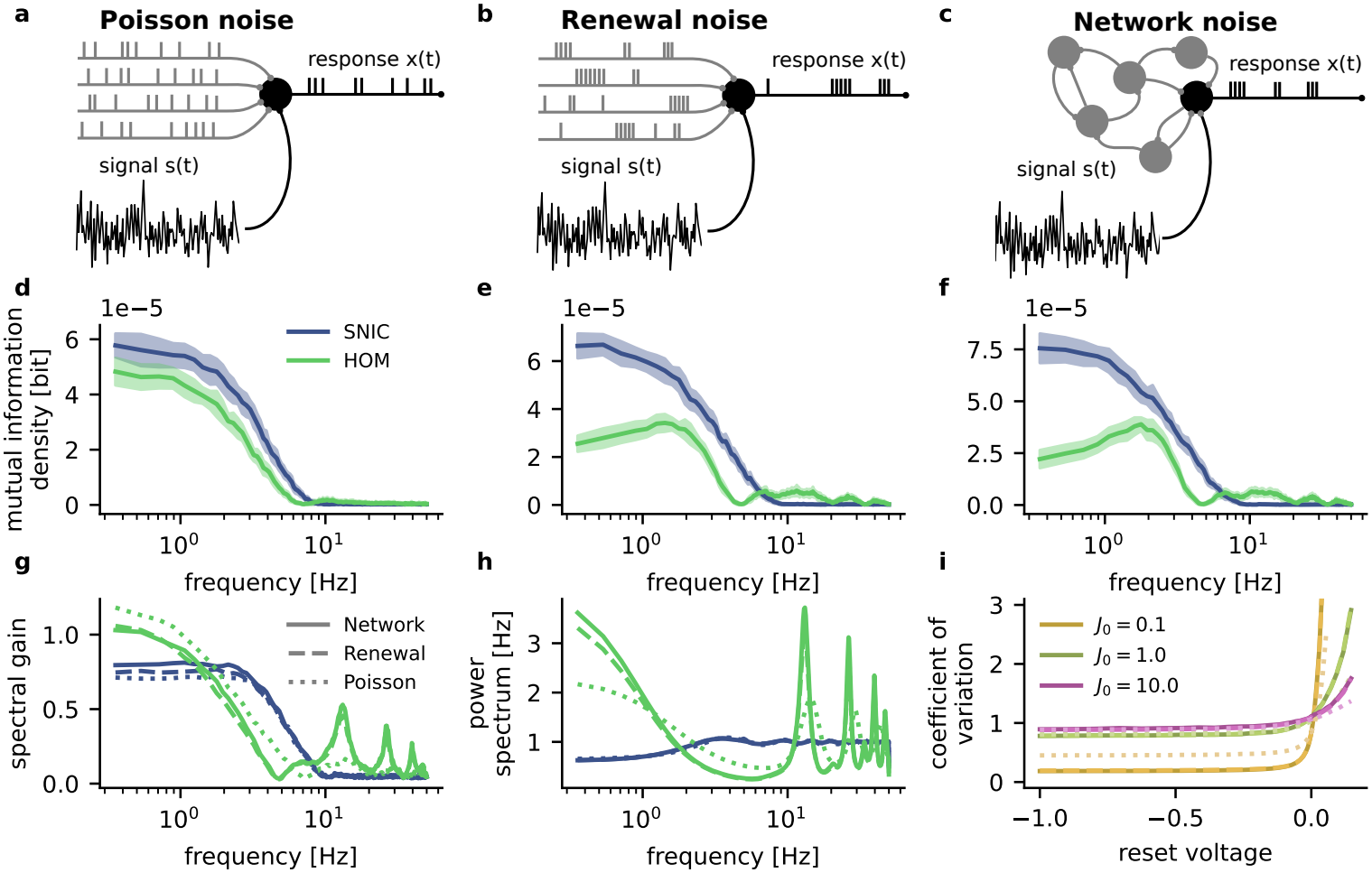
Stimulus encoding in the neural response is compared across three scenarios. In addition to the stimulus **a)** the neuron receives *K* = 20 Poisson inputs as background noise; **b)** the neuron receives *K* = 20 inputs, sampled from the renewal ISI distribution of the network; **c)** the neuron is part of a network and as such receives *K* = 20 recurrent inputs as background noise. **d)** The frequency-resolved mutual information of the HOM neuron (green), compared to the SNIC neuron (blue), is shifted towards intermediate frequencies. This effect is amplified in the network **(f)**. The quantitative match of the renewal approximation **(e)** to the network simulation attributes the amplification to the recurrent input. **g)** The spectral gain of the hom network shows a strong response around the intra-burst frequency. This increased responsiveness is again amplified in the network (solid) and renewal approximation (dashed), compared to the Poisson-driven neuron (dotted). **h)** While a single HOM neuron driven by Poisson input already has increased power in slow frequencies, the self-consistent input in the network and renewal approximation increases this effect. **i)** The CV of the Poisson-driven single neuron qualitatively mirrors the network behavior around the SNL point, while the renewal approximation and network simulation show quantitative agreement over the full range. Notably, for weaker coupling the transition at the SNL point becomes more drastic (see different colors). The simulated networks consist of *N* = 500 neurons, each with *K* = 20 inputs of strength *J*_0_ = 1. The reset is *v*_r_ = *−*0.8 for neurons in the SNIC network and *v*_r_ = 0.115 for HOM. The network activity is tuned to *ν* = 1 Hz.

From a coding perspective, we find that qualitative differences in single-neuron dynamics, induced by the codimension-2 bifurcation at the SNL point, do not wash out but remain present even when the network’s input statistics are shaped self-consistently by its own output — the very regime in which single-cell features were expected to average away.

### Enhanced chaos in the HOM network

The computational capabilities of neural networks rely not only on signal encoding but also on their ability to store and retrieve information, separate inputs, and transform them nonlinearly. An optimal regime for such computation is often located at the edge of chaos [49, 50], and chaotic dynamics themselves have been shown to provide a rich substrate for networks from which they can be trained to perform complex tasks [13]. Quantifying how the excitability switch affects chaos therefore requires isolating its contribution from other sources. This is relevant because excitatory recurrence is itself chaos-promoting: chaos is less intense in inhibitory-only networks than in E-I networks [18]. In fact, substituting inhibitory with excitatory connections, while remaining in the balanced state, can switch a network from a stable to a chaotic state [51]. Restricting this analysis to the purely inhibitory network therefore removes the chaos that E-I structure would generate on its own, so that any increase we observe can be attributed to the critical transition in single-neuron dynamics rather than to the excitatory coupling. We quantify the chaotic network dynamics by calculating the full Lyapunov spectrum of the collective activity (Fig. 3a) through the use of exact event-based network simulations [18]. Lyapunov exponents measure the average rate at which small perturbations of the microscopic network state grow or decay over time. Larger exponents indicate that infinitesimal differences in network state grow faster, making the dynamics less predictable and bearing on the stability and forgetting of memory states [52]. Our analysis shows that the highly irregular dynamics of the HOM network amplify the divergence along unstable directions of the network while increasing the stability otherwise. Overall chaoticity of the network, as captured by the largest Lyapunov exponent, is increased in the HOM regime (Fig. 3b). We explain this effect by the slower time-scales of the burst-like firing, where the unstable directions corresponding to spiking activity change less frequently at the switching between attractors, allowing for stronger divergence.

**Figure 3.**
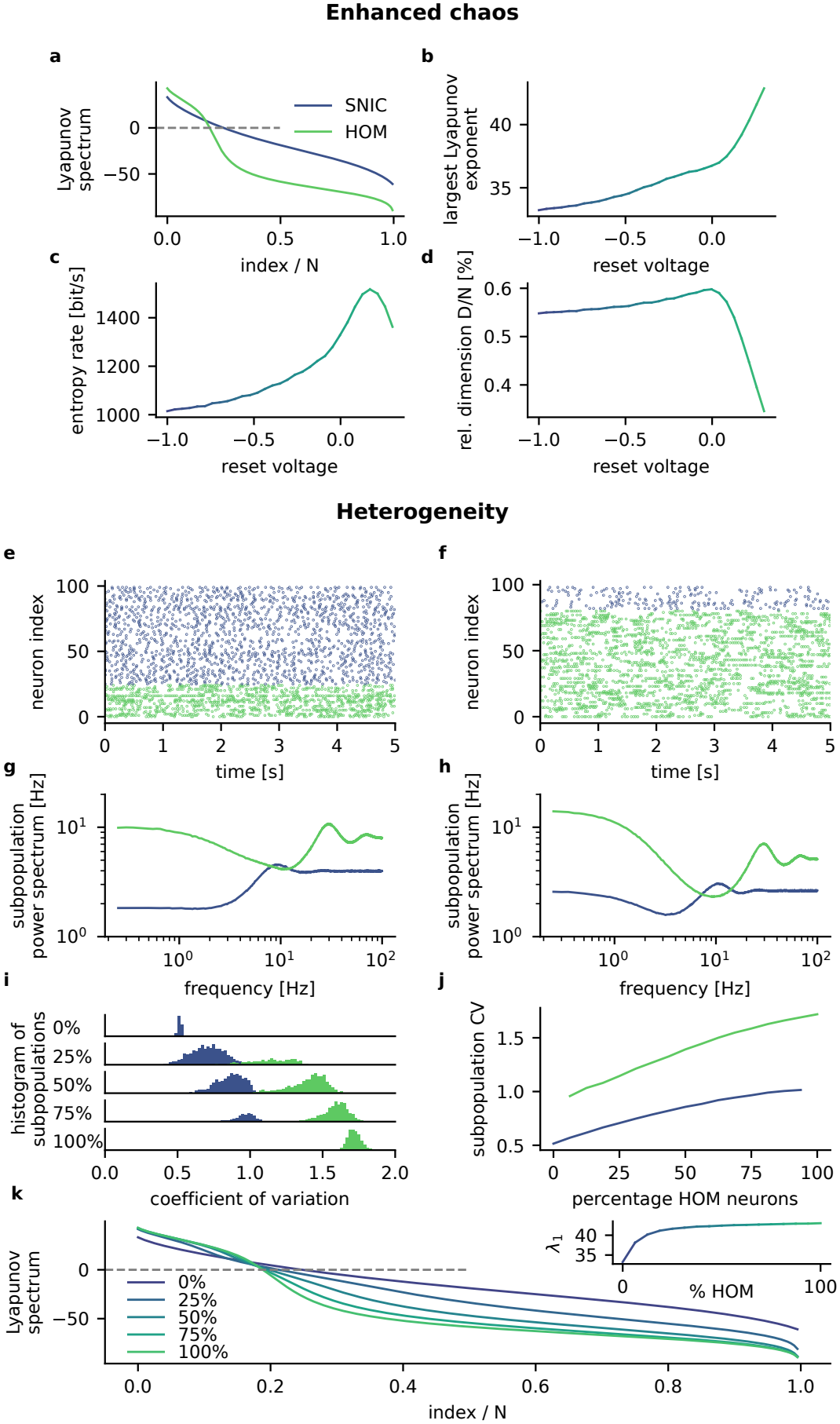
Enhanced network chaos through single cell excitability in networks of heterogeneous onset bifurcations. **a)** Lyapunov spectrum of the recurrent network dynamics of QIF neurons in the SNIC and HOM regimes. **b)** The largest Lyapunov exponent increases in the HOM regime. **c)** The Kolmogorov-Sinai entropy rate, estimated from the sum of positive Lyapunov exponents, peaks in the HOM regime. **d)** The Kaplan-Yorke attractor dimension, based on the number of Lyapunov exponents summing to zero, peaks near the SNL point and decreases in the HOM regime, suggesting a lower-dimensional but more chaotic manifold. **e)** Spike raster plots of a heterogeneous network with 75% SNIC and 25% HOM neurons **(f)** Spike raster of a network with 25% SNIC and 75% HOM neurons. Spiking activity of the subpopulations is largely unaffected by the mixed network. **g)** Power spectra of the SNIC and HOM subpopulations for a network with 75% SNIC and 25% HOM neurons. **h)** Power spectra of the SNIC and HOM subpopulations for a network with 25% SNIC and 75% HOM neurons. **i)** Distributions of single-neuron CVs in the SNIC and HOM subpopulations for 0%, 25%, 50%, 75% and 100% HOM neurons in the network, relative to the number of SNIC neurons. **j)** CV of the subpopulations for varying relative numbers of HOM neurons. **k)** Lyapunov spectra for networks with different relative numbers of HOM neurons. The increase in the largest Lyapunov exponent saturates for relatively small numbers of HOM neurons (see inset). The simulated networks consist of *N* = 200 neurons for the calculation of the Lyapunov spectra, and *N* = 2000 otherwise. Each neuron has *K* = 50 inputs of strength *J*_0_ = 1. The reset is *v*_r_ = *−*0.8 for SNIC neurons and *v*_r_ = 0.115 for HOM. The network activity is tuned to *ν* = 5 Hz.

From the full Lyapunov spectrum, we further calculate the Kolmogorov-Sinai entropy rate, estimated from the sum of the positive Lyapunov exponents, and the Kaplan-Yorke attractor dimension, based on the number of Lyapunov exponents summing to zero [18, 53]. The latter estimates how many degrees of freedom the collective dynamics actively explore: although the network state lives in an N-dimensional space, its trajectory settles onto an attractor of lower dimension, and the Kaplan-Yorke dimension quantifies the size of that subset. We observe an increase in the entropy rate for HOM dynamics, implying more temporal complexity of the network dynamics and stronger sensitivity to microscopic state perturbations (Fig. 3c). The Kaplan-Yorke dimension density D/N is reduced, while the attractor remains extensive (Fig. 3d). Extensivity here means that the attractor dimension still grows in proportion to the size of the network: if the number of neurons is doubled, the number of active dynamical degrees of freedom roughly doubles too. A reduced density D/N signifies that this proportion shrinks in the HOM regime; each neuron contributes less to the collective dynamics, so the trajectory is confined to a smaller fraction of the available dimensions. The chaos thus becomes more intense, as the entropy rate shows, yet unfolds within a lower-dimensional subspace. Notably, both measures (D/N and entropy rate) peak close to the SNL point, in the HOM regime, suggesting a candidate operating point at which the network combines rich internal dynamics with effective exploration of phase space. This observation is not caused by finite size effects as the chaos is extensive (Suppl. Fig. S2).

### Heterogeneous networks

Recent studies have highlighted the importance of heterogeneity in enhancing computational capabilites and robustness in neural circuits [54, 55], where physiological properties vary significantly not only between but also within cell types [56]. To ensure that our key findings are robust in a heterogeneous setting, we introduce heterogeneity in our network model by systematically varying the fraction of neurons in the SNIC and HOM regime. We therefore started with a network comprised exclusively of SNIC neurons and tuned a fraction of randomly chosen neurons into the HOM regime. We incrementally increased the fraction of HOM neurons from 0% to 100% in an otherwise identical network. We then analyzed the resulting network dynamics by calculating the distribution of CVs within the network neurons, as well as the Lyapunov spectra of these heterogeneous networks. Qualitatively, the SNIC and HOM subpopulations exhibit the same firing activity as in homogeneous networks (Fig. 3e, f), showing that neurons retain their excitability class even when they are randomly embedded in a network of neurons with a differing class. With a higher relative number of HOM neurons, reflected in the more variable shared recurrent input, both the SNIC and HOM neurons show increased power in slow frequencies (Fig. 3g, h). The random coupling between the sub-populations lets the distributions of their relative CVs converge (Fig. 3i), while the variablity increases overall (Fig. 3j). Our results further reveal that even a small fraction of HOM neurons drastically affects the network dynamics. As shown in

Fig. 3k, the Lyapunov spectra transition smoothly between the SNIC and the HOM regime as the percentages of HOM neurons increases. Remarkably, the largest Lyapunov exponent *λ*_1_ increases strongly already when only about 25% of the neurons are HOM (see inset). This indicates that a small fraction of HOM neurons can substantially enhance network chaos and variability.

### Mixed Excitatory-inhibitory networks

Since the dynamic interplay of excitation and inhibition is a fundamental organizational principle of cortical networks, we further examine whether the burst-like spiking activity of HOM neurons still persists in excitatory-inhibitory (EI) balanced networks. We find that our main result does extend to this more biologically accurate model, where balance is dynamically maintained by an additional excitatory population, corroborating the general validity of our results with the prototypical model of asynchronous irregular activity in the brain [1, 2, 31, 33].

In the same manner as the purely inhibitory networks previously discussed, we observe the Poisson-like asynchronous irregular firing of the EI network with SNIC onset (Fig. 4a) to become even more variable in the HOM regime (Fig. 4b), with CVs surpassing one around the SNL point (Fig. 4d). The ISI distribution of the HOM network shows the most prominent difference introduced by the excitatory population, with a sharp peak at the intrinsic limit-cycle period being broadened. In contrast to the purely inhibitory network, the recurrent excitation in the EI network can not only cause delays but also advances in the phase of a neuron, leading to faster spiking. Because bistable dynamics are often associated with interneurons, we further examine a network in which only the inhibitory population has HOM dynamics (Fig. 4e), demonstrating cross-talk between the populations in a similar manner to the heterogeneous network previously discussed (compare Fig. 4f and Fig. 3i), highlighting that only the small inhibitory subset of HOM neurons can alter collective behavior of the entire network.

**Figure 4.**
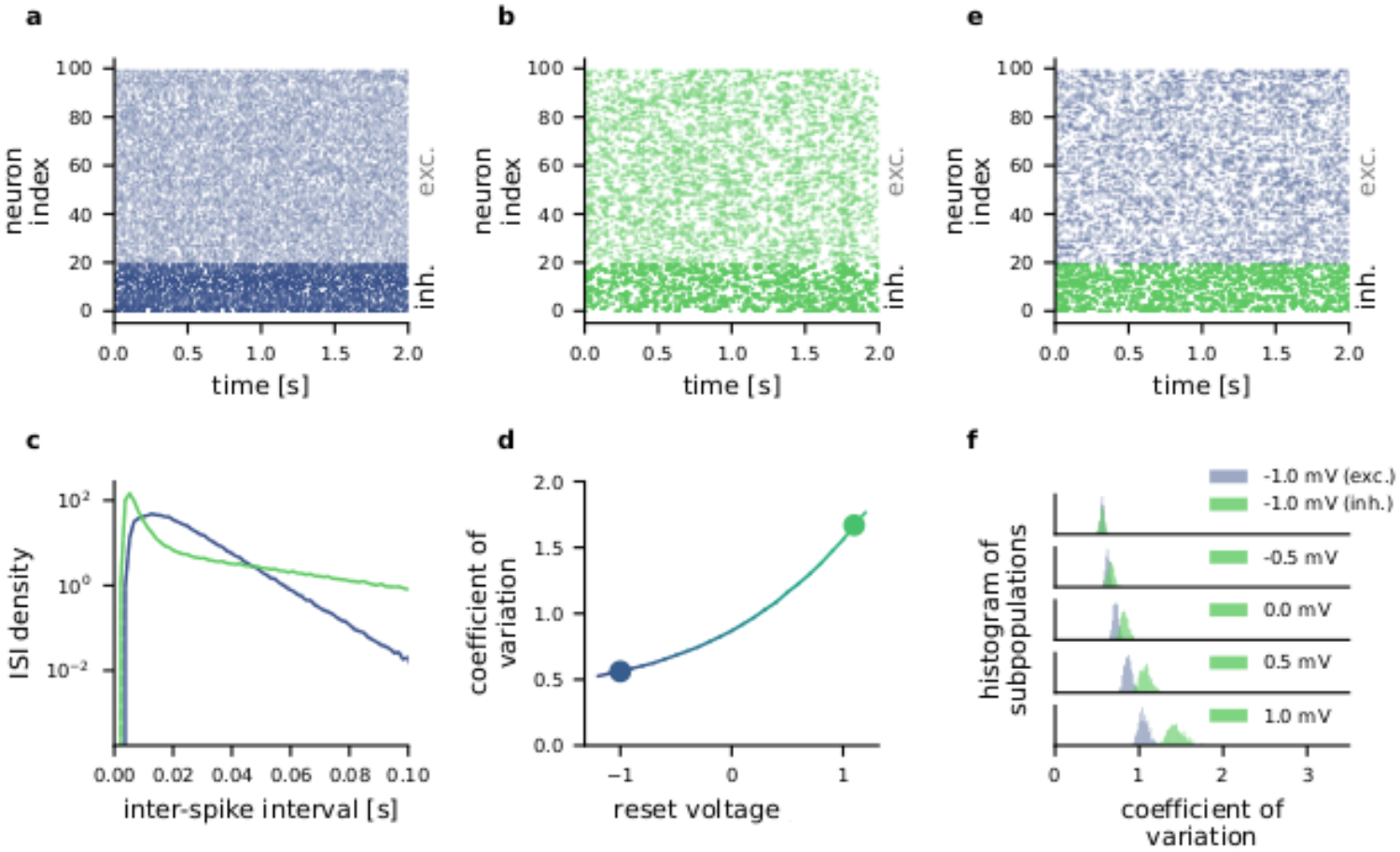
Excitatory-inhibitory balanced networks with SNIC and HOM excitability. **a)** Spike raster of an EI network of SNIC neurons with Poisson-like activity. **b)** Spike raster of the same network with HOM excitability exhibits burst-like firing. **c)** Densities of inter-spike intervals for EI networks with SNIC and HOM excitability. **d)** The coefficient of variation of the inter-spike intervals increases gradually with the reset voltage, switching from sub-Poissonian (SNIC) to super-Poissonian (HOM) at the SNL point. **e)** Spike raster of an EI network where the burst-like activity of inhibitory HOM neurons affects excitatory SNIC neurons. **f)** Histograms of CVs of the inhibitory and excitatory subpopulations for increasing reset voltages of the inhibitory neurons. The bottom panel corresponds to network in **(e)**, demonstrating the cross-population influence of HOM dynamics. The network model consists of *N*_E_ = 1600 excitatory and *N*_I_ = 400 inhibitory neurons. Each neuron receives *K*_I_ = 30 inhibitory and *K*_E_ = 120 excitatory inputs. Coupling strength is *J*_E_ = 1 for excitatory and *J*_I_ = *−*5 for inhibitory synapses . The reset is *v*_r_ = *−*1.0 for SNIC neurons and *v*_r_ = 1.0 for HOM, if not indicated otherwise. The network activity is tuned to *ν* = 20 Hz.

Since in addition to HOM dynamics, interneurons are also known to exhibit a Hopf spike-onset [57, 58], we provide a similar analysis of networks with bistable Hopf neurons in Fig. S4 and Fig. S5. These simulations show that networks of Hopf neurons in their bistable regime also exhibit highly variable activity with burst-like firing of individual units, consistent with our main claim that single-neuron bifurcation properties translate directly to the collective dynamics of the network. Thus, the central conclusion is supported for both HOM and Hopf spike onsets.

## I. DISCUSSION

Computations in cortical networks are generally believed to benefit from rich internal dynamics, resulting from appropriately tuned network properties. Highly variable spiking activity in particular is often explained by the topology and strength of connections as well as synaptic timescales. Here, we demonstrate a novel mechanism, where a network state transition – marked by a steep increase in spiking irregularity – originates from a critical switch in single neuron dynamics. It is founded in the very process of action potential generation described by the mathematical bifurcation underlying the spike onset. Specifically, by inducing a ubiquitous biological switch in the excitability of neurons, we show that the highly variable burst-like activity associated with a homoclinic spike onset not only persists in a large balanced network, but is even amplified by the recurrent interactions. The singleneuron nature of this mechanism is supported by a self-consistent renewal approximation over a large range of parameters. We demonstrate the influence of homoclinic neuronal dynamics on the collective network activity to shift information transfer from lower to intermediate and higher frequencies while also enhancing chaos on a lower-dimensional attractor, highlighting the profound impact of single-neuron physiology on the computational capabilities of balanced networks. The effects on spike variability also persist in networks with multiple populations of either excitatory and inhibitory neurons or different excitability classes, and do not rely on a particular choice of neuron model, showing the general validity of our findings.

### Relation to previous work

Our results qualify the view that in the asynchronous regime single-neuron properties can be absorbed into the gain function and connectivity strength [33, 59], and are more in line with work linking single-cell and connectivity detail to collective dynamics [60, 61]. The asynchronous irregular firing observed in these networks has gathered considerable attention as a model for cortical activity. Particularly dynamic regimes with even higher variability have so far been attributed to synaptic properties [9–11, 62, 63]. Contrary to this view, we demonstrate that intrinsic neuronal features, specifically the excitability class, can substantially shape the state of a network, leading to similar collective dynamics without changing the connections.

While previous studies examining the role of HOM excitability focused either on the level of single neurons [20, 22, 29], or on the context of specified small networks [23, 64], our work extends the scope of these previous findings to the fluctuation driven regime of larger random networks caused by the balance of excitation and inhibition.

We emphasize that the parameter changes that switch SNIC to HOM excitability induce a codim-2 SNL bifurcation with a significant qualitative change in single neuron dynamics and a profound impact on network properties. While changes in other parameters or heterogeneous distributions of those parameters have also been studied [34, 35], these variations often do not correspond to a critical transition in the underlying single-neuron dynamics, leading to quantitative effects rather than qualitatively different network behavior. A complementary effect to the HOM excitability in QIF neurons, where the increased reset voltage corresponds to a faster downstroke of the action potential, is presented in a study about the rapidness of the upstroke [65]. While also impacting information filtering and chaotic dynamics on the network level, it relates to the biophysics of sodium, rather than potassium channels and does not cause a change in the excitability class. Yet other studies have examined the impact of the phase-response curve of a neuron, which can be associated with its excitability class, on the network level [66, 67], yet not in the fluctuation-driven balanced regime examined here.

The results presented in this study therefore fill an important gap by examining the impact of a physiological property of neurons that is of particular interest for the qualitative dynamic changes it induces, studied in the context of the large random networks believed to be prevalent in the mammalian cortex.

### Relevance and Implications

#### Generality

The examined excitability switch is far from an idiosyncrasy of the model we present. HOM excitability can be induced in a vast class of neurons [28] via biologically relevant changes such as variations in temperature [22], extracellular potassium concentration [29], morphology [30], and ion channel composition [23]. Collectively, these findings suggest that this mechanism is likely to be a ubiquitous and functionally significant feature in the brain.

#### Neurobiological relevance

The highly irregular burst-like network dynamics that we observe, together with the reported changes in information transfer and enhanced chaotic dynamics could have implications for both the computational capabilities as well as pathological behavior of cortical networks. In recent years, there has been considerable interest in bursts as a ternary neuronal code, complementing the binary code of silence and isolated spikes [68]. Short intervals of high-frequency firing could impact plasticity and learning in a way that is distinct from overall increases of the firing rate. Further, bursting has been shown to enable multiplexed coding of information [69] and has even been proposed as a biological implementation of backpropagation of error signals [70, 71]. These mechanisms rely on deterministically bursting neurons, but specific input could induce deterministic bursts in the bistable HOM dynamics as well. While in general, this kind of bursting is attributed to calcium dynamics, our model points to a different mechanism which could help understand burst-like dynamics in superficial cortical layers where the required calcium spiking is not observed experimentally [72]. However, homoclinic spiking might also be linked to certain neurological conditions characterized by abnormal network dynamics, such as febrile seizures associated with a rise in temperature [22]. Accumulation of extracellular potassium, which is also associated with bursting and overall increased activity during epileptic seizures [73, 74], could relate to the induced HOM excitability [29].

The observed shift in information encoding to intermediate frequencies suggests an improved selective response of the network to specific input signals. This band-pass filtering effect could allow networks with HOM excitability to encode and process relevant information at distinct frequencies while suppressing slow noise and drifts, thus improving overall computational efficiency [75].

The peak in entropy production and attractor dimension near the SNL point (Fig. 3c,d) is suggestive in a computational context: the rich internal dynamics of this network state could mark a favourable operating point for subsequent training [13]. The effect is twofold — the dynamics become more chaotic overall, yet remain confined to a lower-dimensional attractor, combining sensitivity to inputs with a compact representation. Notably, where such dynamics originate also determines how they can be controlled: if high variability arises from connectivity or synaptic strength, tuning it necessarily acts on the same parameters that carry learning. A cellular mechanism separates the two, allowing a network’s dynamical regime to be set independently of its synaptic weights.

Our results link macroscopic network behavior to the physiological properties of single neurons, highlighting the importance of intrinsic neuronal dynamics for both complex computations as well as pathological behavior of neural circuits.

### Limitations of the analysis

While our model captures the core mechanics of both HOM excitability and the balance of excitation and inhibition in cortical networks, it relies on several simplifications that warrant further examination. The use of pulse-coupled QIF neurons offers a model of HOM network dynamics that is both analytically tractable and computationally efficient. This approach is justified near the excitability switch, where the QIF model represents the unfolding of the SNL bifurcation, but changes from the observed behavior are expected for significant deviations from this operating point. While we observe qualitatively consistent results with conductance-based models in this regime, biological mechanisms that induce a transition of the excitability switch can also introduce secondary effects, further altering the bifurcation structure [76].

In addition, the simplified treatment of synaptic interactions as instantaneous pulses, used here to separate the single-neuron excitability from synaptic effects, neglects potentially important temporal dynamics. Although the influence of HOM excitability decreases for larger synaptic time constants, its effects remain significant within biologically relevant ranges (e.g. GABA_A_ *≈*5 ms) [22].

Addressing these limitations in future work will require the integration of more complex neuron models with dynamic synapses in biologically realistic network architectures.

The information theoretic analysis presented here examines the information of individual neurons in the network, instead of postulating that information is present in the mean population activity. Both approaches neglect information that is stored in the correlative structure of population activities. Given that the emergence of irregular firing in the present article is not tied to strong network weights, correlations are assumed to be weak and are, therefore, neglected.

### Outlook

The rich collective dynamics caused by the HOM excitability suggest an interesting line of research in the context of task-specific artificial spiking neural networks. In the case of liquid state machines, the optimal operating point of these networks is widely regarded to be on the edge of chaos [49, 50, 77], where separability of inputs is balanced with fading memory timescales. While the balanced network we studied is highly chaotic, the mechanism of the excitability switch could allow the dynamics of different networks operating in ordered regimes to be tuned towards chaos without interfering with the synaptic parameters required for most learning algorithms. While to date it remains an open question to what extent chaos is beneficial as a starting point for learning [78], the presented rich chaotic dynamics near the SNL point could present a substrate for complex computations as a basis for FORCE learning [13] or similar approaches. The slow timescales of the burst-like activity could be particularly useful for temporal tasks, like the generation of slowly varying signals, without the need for additional adaptation currents. We therefore argue that the QIF neuron model presents a computationally efficient alternative to the prevalent leaky integrate-and-fire neuron that can enhance network dynamics substantially.

Overall, our results show that HOM excitability at the single-neuron level can profoundly shape collective network dynamics, giving rise to a state of highly variable, burst-like spiking activity, altering information encoding, and enhancing chaos. These findings bridge the gap between cellular biophysics and macroscopic neural computations. They identify the conditions under which intrinsic neuronal properties escape absorption by the collective state, and thereby acquire direct relevance for cortical network dynamics.

## Methods

### Phase description of the QIF neuron model

The quadratic integrate and fire (QIF) is a simple neuron model that nonetheless captures the transition between spike onset-bifurcations that we will investigate here. It is governed by a single equation

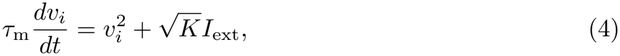

with a constant external bias current *I*_ext_ and a membrane time constant of *τ*_m_ = 10 ms used throughout the article. When the membrane voltage diverges to infinity *v → ∞* in finite time, it is reset to a fixed voltage *v*_r_. In the common configuration with the reset at *v*_r_ = *→ ∞*, the QIF neuron is the normal form of the SNIC spike onset-bifurcation which can be found in all conductance-based neuron models. We use the reset voltage as a bifurcation parameter to induce the switch between SNIC and HOM onset-bifurcation by increasing it past the SNL codimension two SNL point at *v*_r_ = 0. The QIF model can be integrated analytically. For example for *v*_r_ = *→ ∞*, the time evolution of the membrane voltage between spikes is given by

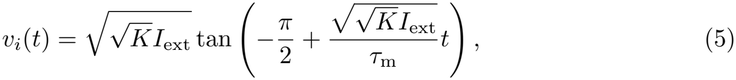

showing the convex spike shape and divergence in finite time. The analytical solution allows an exact phase representation by change of variables [38]. Substituting 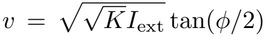 leads to a phase representation with constant phase velocity inbetween incoming spikes

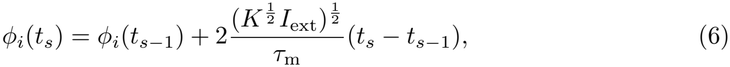

with reset phase *ϕ*_*r*_ = 2 arctan 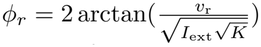 and spike threshold at *ϕ*_*th*_ = *π*. In the case of pulse input, all the response of the neuron can be captured exactly through the phase transition curve (PTC), which is readily derived from the phase representation as

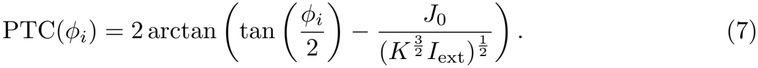

### Network model and simulations

The network model considered here is a sparse, pulse-coupled, inhibitory balanced network of quadratic integrate and fire (QIF) neurons, adapted from previous studies on spiking balanced networks [18]. Unless otherwise stated in the corresponding figure captions, the network consists of *N* = 1000 QIF neurons, each receiving inputs from a fixed number of *K* = 30 randomly drawn presynaptic neurons in the network. The recurrent connectivity is purely inhibitory with a homogeneous coupling strength *J*_0_ = 1 which is scaled by the in-degree 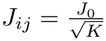. Accordingly, the recurrent input consists of a sum over delta functions at the presynaptic spike times, resulting in the sub-threshold dynamics

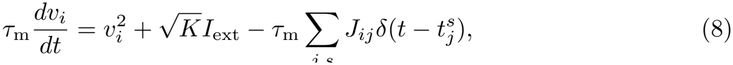

a term for the sum over all recurrent network inputs is added to Eq. (4). Because HOM neurons are more active, the constant bias input current *I*_ext_ is adjusted to tune the network firing rate to a predefined value *ν* = 1 Hz. The only sources of randomness in the network are the quenched noise of the network connectivity and initial conditions. Without any external noise, the system is fully deterministic.

For the simulation of the network, two methods are used. Some results are produced with fixed step-size integration of the network equations. Other simulations are based on the phase description of the QIF model. The linear phase evolution between spikes allows a precise event-based simulation of the network, where only the interactions at spike time need to be computed and the time to the next spike has to be kept track of. For very sparse networks, this event-based simulation method offers a considerable reduction in computational cost while being exact to numerical precision [79]. While we show, that the main results of this study do not rely on the simulation method, the event-based simulation primarily allows the calculation of the Lyapunov spectra. Based on the phase description of the network, the interactions during spike time can be used to derive a single-spike Jacobian from the PTC [18]. Using this Jacobian, the Lyapunov spectra are calculated in the standard Gram-Schmidt re-orthonormalization procedure. From the Lyapunov exponents *λ*_*i*_, the Kolmogorov-Sinai entropy production is calculated via the Pesin identity as 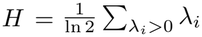 and the Kaplan-Yorke attractor dimension as 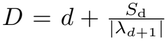 where *d* is the largest number of exponents so that 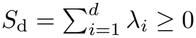.

### Renewal Approximation

We use a renewal approximation developed in previous studies [46, 80, 81] as a numerical mean-field theory, to self-consistently compute the firing statistics of the network. While a self-consistent mean-field description can be derived analytically for a network of SNIC neurons, the required diffusion approximation of the inputs is not valid in the case of a HOM network, where the recurrent input is colored by the slow burst-like network activity.

This renewal approach relies on recording the ISI distribution of a single neuron receiving an increasingly better approximation of network input through an iterative approach. The neuron is initially driven by Poisson spikes and its response is recorded. The ISI distribution of the response is then sampled to create new surrogate input in the next iteration and the output is recorded again. These steps are repeated until input and output of the neuron are statistically equivalent. Convergence is monitored via the firing rate and the CV of the ISI distribution.

Three situations are compared. A neuron receiving Poisson input, a neuron receiving renewal input and a neuron receiving recurrent input in the network. Input rates in the Poisson and renewal case are adjusted to match the statistics in the network, meaning the same number *K* of inputs with the same mean firing rate *ν*, which the network is tuned to.

### Information theory

To characterize how the computational capabilities of the balanced network are affected by the switch in excitability, we employ Shannon’s information theory as an agnostic, model-free approach. We compare the amount of information which is encoded by a network of SNIC and HOM neurons. To analyze filtering properties, rather than overall amount of information, we approximate a frequency-resolved measure, a density of Shannon’s mutual information rate (MIR). For a Gaussian input signal, a lower bound of the MIR can be derived from the squared spectral coherence [82]

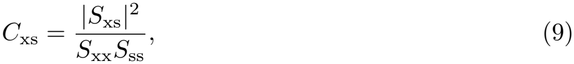

which captures frequency-dependent correlations between the input signal and network response. Here, *S*_ss_ and *S*_xx_ are the auto spectral densities of the signal and the response. The cross spectral density *S*_xs_ is also used to calculate the gain *G* = *S*_xs_*/S*_ss_. The direct calculation of the mutual information rate requires a large amount of data, it is not feasible here. Yet, the lower bound for the mutual information rate

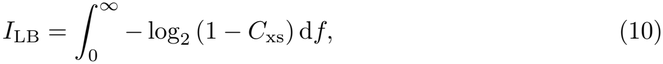

has been shown to be a tight bound in related cases [44]. We focus here not on the total amount of information transfer but rather on the frequency-dependent MIR density given by the integrand

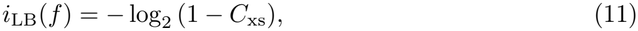

which quantifies information encoding resolved over frequency bands.

The signal is a high-frequency shot noise approximation of Gaussian white noise. It is implemented as a sum of an excitatory and an inhibitory Poisson input with a rate of *ν*_s_ = 10 kHz each, necessitated by the event-based simulation. We verify that the signal is sufficiently weak to measure a linear response (see Supplement Fig. S1).

## ACKNOWLEDGEMENTS

This project has received funding from the European Research Council (ERC) under the European Union’s Horizon 2020 research and innovation program (grant number 864243), from the German Research Council (CRC/TRR 384), and the Einstein Foundation Berlin (grant number EP-2021-621).

## References

[1] N. Brunel, Dynamics of Sparsely Connected Networks of Excitatory and Inhibitory Spiking Neurons, Journal of Computational Neuroscience 8, 183 (2000).

[2] C. van Vreeswijk and H. Sompolinsky, Chaotic Balanced State in a Model of Cortical Circuits, Neural Computation 10, 1321 (1998).

[3] A. Compte, C. Constantinidis, J. Tegnér, S. Raghavachari, M. V. Chafee, P. S. Goldman-Rakic, and X.-J. Wang, Temporally Irregular Mnemonic Persistent Activity in Prefrontal Neurons of Monkeys During a Delayed Response Task, Journal of Neurophysiology 90, 3441 (2003).

[4] M. Shafi, Y. Zhou, J. Quintana, C. Chow, J. Fuster, and M. Bodner, Variability in neuronal activity in primate cortex during working memory tasks, Neuroscience 146, 1082 (2007).

[5] E. D. Gershon, M. C. Wiener, P. E. Latham, and B. J. Richmond, Coding Strategies in Monkey V1 and Inferior Temporal Cortices, Journal of Neurophysiology 79, 1135 (1998).

[6] S. Shinomoto, K. Shima, and J. Tanji, Differences in spiking patterns among cortical neurons, Neural computation 15, 2823 (2003).

[7] G. Maimon and J. A. Assad, Beyond Poisson: Increased spike-time regularity across primate parietal cortex, Neuron 62, 426 (2009).

[8] H. Sompolinsky, A. Crisanti, and H. J. Sommers, Chaos in Random Neural Networks, Physical Review Letters 61, 259 (1988).

[9] A. Lerchner, C. Ursta, J. Hertz, M. Ahmadi, P. Ruffiot, and S. Enemark, Response variability in balanced cortical networks, Neural computation 18, 634 (2006).

[10] S. Ostojic, Two types of asynchronous activity in networks of excitatory and inhibitory spiking neurons, Nature Neuroscience 17, 594 (2014).

[11] A. Litwin-Kumar and B. Doiron, Slow dynamics and high variability in balanced cortical networks with clustered connections, Nature Neuroscience 15, 1498 (2012).

[12] D. Hansel and G. Mato, Asynchronous states and the emergence of synchrony in large networks of interacting excitatory and inhibitory neurons, Neural computation 15, 1 (2003).

[13] D. Sussillo and L. F. Abbott, Generating coherent patterns of activity from chaotic neural networks, Neuron 63, 544 (2009).

[14] M. Rabinovich, R. Huerta, and G. Laurent, Transient Dynamics for Neural Processing, Science 321, 48 (2008).

[15] E. M. Izhikevich, Dynamical Systems in Neuroscience (MIT press, 2007).

[16] A. Palmigiano, R. Engelken, and F. Wolf, Boosting of neural circuit chaos at the onset of collective oscillations, eLife 12 (2023).

[17] S. Jahnke, R.-M. Memmesheimer, and M. Timme, Stable Irregular Dynamics in Complex Neural Networks, Physical Review Letters 100, 048102 (2008).

[18] M. Monteforte and F. Wolf, Dynamical Entropy Production in Spiking Neuron Networks in the Balanced State Supplementary Material, Physical Review Letters 105, 268104 (2010).

[19] D. Durstewitz and T. Gabriel, Dynamical basis of irregular spiking in NMDA-driven prefrontal cortex neurons, Cerebral cortex 17, 894 (2007).

[20] J.-H. Schleimer, J. Hesse, S. A. Contreras, and S. Schreiber, Firing statistics in the bistable regime of neurons with homoclinic spike generation, Physical Review E 103, 012407 (2021).

[21] J. Hesse, J.-H. Schleimer, and S. Schreiber, Qualitative changes in phase-response curve and synchronization at the saddle-node-loop bifurcation, Physical Review E 95, 052203 (2017).

[22] J. Hesse, J.-H. Schleimer, N. Maier, D. Schmitz, and S. Schreiber, Temperature elevations can induce switches to homoclinic action potentials that alter neural encoding and synchronization, Nature Communications 13, 3934 (2022).

[23] S. Hürkey, N. Niemeyer, J.-H. Schleimer, S. Ryglewski, S. Schreiber, and C. Duch, Gap junctions desynchronize a neural circuit to stabilize insect flight, Nature 618, 118 (2023).

[24] A. Arvanitaki, Les variations graduées de la polarisation des systèmes excitables: Relation avec la négativité propagée et signification fonctionnelle dans l’activité rhythmique, (1938).

[25] A. L. Hodgkin, The local electric changes associated with repetitive action in a non-medullated axon, The Journal of physiology 107, 165 (1948).

[26] J. Rinzel and G. B. Ermentrout, Analysis of neural excitability and oscillations, Methods in neuronal modeling 2, 251 (1998).

[27] J.-H. Schleimer and S. Schreiber, Phase-response curves of ion channel gating kinetics, Mathematical Methods in the Applied Sciences 41, 8844 (2018).

[28] C. Kirst, J. Ammer, F. Felmy, A. Herz, and M. Stemmler, Fundamental structure and modulation of neuronal excitability: Synaptic control of coding, resonance, and network synchronization, BioRxiv, 022475 (2015).

[29] S. A. Contreras, J.-H. Schleimer, A. T. Gulledge, and S. Schreiber, Activity-mediated accumulation of potassium induces a switch in firing pattern and neuronal excitability type, PLoS Computational Biology 17, e1008510 (2021).

[30] R. P. Gowers and S. Schreiber, How neuronal morphology impacts the synchronisation state of neuronal networks, PLOS Computational Biology 20, e1011874 (2024).

[31] A. Renart, J. de la Rocha, P. Bartho, L. Hollender, N. Parga, A. Reyes, and K. D. Harris, The Asynchronous State in Cortical Circuits, Science 327, 587 (2010).

[32] D. J. Amit and N. Brunel, Dynamics of a recurrent network of spiking neurons before and following learning, Network: Computation in Neural Systems 8, 373 (1997).

[33] C. Van Vreeswijk and H. Sompolinsky, Chaos in Neuronal Networks with Balanced Excitatory and Inhibitory Activity, Science 274, 1724 (1996).

[34] J. F. Mejias and A. Longtin, Differential effects of excitatory and inhibitory heterogeneity on the gain and asynchronous state of sparse cortical networks, Frontiers in computational neuroscience 8, 107 (2014).

[35] S. Khajehabdollahi, R. Zeraati, E. Giannakakis, T. J. Schäfer, G. Martius, and A. Levina, Emergent mechanisms for long timescales depend on training curriculum and affect performance in memory tasks (2024), arXiv:2309.12927 [cs].

[36] S. Schecter, The Saddle-Node Separatrix-Loop Bifurcation, SIAM Journal on Mathematical Analysis 18, 1142 (1987).

[37] S.-N. Chow and X.-B. Lin, Bifurcation of a homoclinic orbit with a saddle-node equilibrium, (1990).

[38] B. Ermentrout, Type I membranes, phase resetting curves, and synchrony, Neural computation 8, 979 (1996).

[39] N. Brunel and V. Hakim, Fast global oscillations in networks of integrate-and-fire neurons with low firing rates, Neural computation 11, 1621 (1999).

[40] J.-H. Schleimer and M. Stemmler, Coding of Information in Limit Cycle Oscillators, Physical Review Letters 103, 248105 (2009).

[41] J. Hesse, Implications of neuronal excitability and morphology for spike-based information transmission, (2017).

[42] T. Herfurth and T. Tchumatchenko, Information transmission of mean and variance coding in integrate-and-fire neurons, Physical Review E 99, 032420 (2019).

[43] F. Rieke, D. Warland, R. d. R. Van Steveninck, and W. Bialek, Spikes: Exploring the Neural Code (MIT press, 1999).

[44] D. Bernardi and B. Lindner, A frequency-resolved mutual information rate and its application to neural systems, Journal of Neurophysiology 113, 1342 (2015).

[45] R. D. Vilela and B. Lindner, Comparative study of different integrate-and-fire neurons: Spontaneous activity, dynamical response, and stimulus-induced correlation, Physical Review E 80, 031909 (2009).

[46] B. Dummer, S. Wieland, and B. Lindner, Self-consistent determination of the spike-train power spectrum in a neural network with sparse connectivity, Frontiers in Computational Neuroscience 8, 104 (2014).

[47] M. Di Volo and A. Torcini, Transition from Asynchronous to Oscillatory Dynamics in Balanced Spiking Networks with Instantaneous Synapses, Physical Review Letters 121, 128301 (2018).

[48] M. Di Volo, M. Segneri, D. S. Goldobin, A. Politi, and A. Torcini, Coherent oscillations in balanced neural networks driven by endogenous fluctuations, Chaos: An Interdisciplinary Journal of Nonlinear Science 32 (2022).

[49] R. Legenstein and W. Maass, Edge of chaos and prediction of computational performance for neural circuit models, Neural Networks Echo State Networks and Liquid State Machines, 20, 323 (2007).

[50] N. Bertschinger and T. Natschläger, Real-Time Computation at the Edge of Chaos in Recurrent Neural Networks, Neural Computation 16, 1413 (2004).

[51] S. Jahnke, How chaotic is the balanced state?, Frontiers in Computational Neuroscience 3, 10.3389/neuro.10.013.2009 (2009).

[52] J. Schuecker, S. Goedeke, and M. Helias, Optimal Sequence Memory in Driven Random Networks, Physical Review X 8, 041029 (2018).

[53] Y. B. Pesin, Characteristic Lyapunov exponents and smooth ergodic theory, Russian Mathematical Surveys 32, 55 (1977).

[54] N. Perez-Nieves, V. C. H. Leung, P. L. Dragotti, and D. F. M. Goodman, Neural heterogeneity promotes robust learning, Nature Communications 12, 5791 (2021).

[55] R. Gast, S. A. Solla, and A. Kennedy, Neural heterogeneity controls computations in spiking neural networks, Proceedings of the National Academy of Sciences 121, e2311885121 (2024).

[56] E. Marder and A. L. Taylor, Multiple models to capture the variability in biological neurons and networks, Nature Neuroscience 14, 133 (2011).

[57] D. Golomb, K. Donner, L. Shacham, D. Shlosberg, Y. Amitai, and D. Hansel, Mechanisms of firing patterns in fast-spiking cortical interneurons, PLoS Computational Biology 3, e156 (2007).

[58] J. Ševčík and L. Přibylová, Cycle multistability and synchronization mechanisms in coupled interneurons: In-phase and anti-phase dynamics under current stimuli, Applied Mathematics and Computation 503, 129500 (2025).

[59] W. Gerstner, W. M. Kistler, R. Naud, and L. Paninski, Neuronal Dynamics (Cambridge University Press, 2014).

[60] M. di Volo, A. Romagnoni, C. Capone, and A. Destexhe, Biologically realistic mean-field models of conductance-based networks of spiking neurons with adaptation, Neural Computation 31, 653 (2019).

[61] F. Mastrogiuseppe and S. Ostojic, Linking connectivity, dynamics, and computations in low-rank recurrent neural networks, Neuron 99, 609 (2018).

[62] O. Harish and D. Hansel, Asynchronous rate chaos in spiking neuronal circuits, PLoS computational biology 11, e1004266 (2015).

[63] R. Engelken, F. Farkhooi, D. Hansel, C. van Vreeswijk, and F. Wolf, A reanalysis of “Two types of asynchronous activity in networks of excitatory and inhibitory spiking neurons”, F1000Research 5 (2016).

[64] L. Weerdmeester, N. Niemeyer, P. Pfeiffer, S. Billaudelle, J. Schemmel, J.-H. Schleimer, andS. Schreiber, Qualitative switches in single-neuron spike dynamics on neuromorphic hardware: Implementation, impact on network synchronization and relevance for plasticity, Neuromorphic Computing and Engineering 4, 014009 (2024).

[65] R. Engelken, M. Monteforte, and F. Wolf, Sparse chaos in cortical circuits (2024), arXiv:2412.21188 [q-bio].

[66] J. Ladenbauer, M. Augustin, L. Shiau, and K. Obermayer, Impact of Adaptation Currents on Synchronization of Coupled Exponential Integrate-and-Fire Neurons, PLOS Computational Biology 8, e1002478 (2012).

[67] D. Pazó, E. Montbrió, and R. Gallego, The winfree model with heterogeneous phase-response curves: Analytical results, Journal of Physics A: Mathematical and Theoretical 52, 154001 (2019).

[68] Z. Friedenberger, E. Harkin, K. Tóth, and R. Naud, Silences, spikes and bursts: Three-part knot of the neural code, The Journal of Physiology 601, 5165 (2023).

[69] R. Naud and H. Sprekeler, Sparse bursts optimize information transmission in a multiplexed neural code, Proceedings of the National Academy of Sciences 115, 10.1073/pnas.1720995115 (2018).

[70] A. Payeur, J. Guerguiev, F. Zenke, B. A. Richards, and R. Naud, Burst-dependent synaptic plasticity can coordinate learning in hierarchical circuits, Nature Neuroscience 24, 1010 (2021).

[71] M. Stuck and R. Naud, Burstprop for Learning in Spiking Neuromorphic Hardware, in Proceedings of the 2023 International Conference on Neuromorphic Systems (ACM, Santa Fe NM USA, 2023) pp. 1–5.

[72] M. E. Larkum, J. Waters, B. Sakmann, and F. Helmchen, Dendritic spikes in apical dendrites of neocortical layer 2/3 pyramidal neurons, Journal of Neuroscience 27, 8999 (2007).

[73] S. J. Korn, J. L. Giacchino, N. L. Chamberlin, and R. Dingledine, Epileptiform burst activity induced by potassium in the hippocampus and its regulation by GABA-mediated inhibition, Journal of Neurophysiology 57, 325 (1987).

[74] M. S. Jensen and Y. Yaari, Role of Intrinsic Burst Firing, Potassium Accumulation, and Electrical Coupling in the Elevated Potassium Model of Hippocampal Epilepsy, Journal of Neuro-physiology 77, 1224 (1997).

[75] S. Sinanović and D. H. Johnson, Toward a theory of information processing, signal processing 87, 1326 (2007).

[76] M. Behbood, L. Lemaire, J.-H. Schleimer, and S. Schreiber, The Na+/K+-ATPase generically enables deterministic bursting in class I neurons by shearing the spike-onset bifurcation structure, PLOS Computational Biology 20, e1011751 (2024).

[77] C. G. Langton, Computation at the edge of chaos: Phase transitions and emergent computation, Physica D: Nonlinear Phenomena 42, 12 (1990).

[78] R. Engelken, A. Ingrosso, R. Khajeh, S. Goedeke, and L. F. Abbott, Input correlations impede suppression of chaos and learning in balanced firing-rate networks, PLOS Computational Biology 18, 1 (2022).

[79] R. Engelken, Sparseprop: Efficient event-based simulation and training of sparse recurrent spiking neural networks, Advances in Neural Information Processing Systems 36, 3638 (2023).

[80] R. F. O. Pena, S. Vellmer, D. Bernardi, A. C. Roque, and B. Lindner, Self-Consistent Scheme for Spike-Train Power Spectra in Heterogeneous Sparse Networks, Frontiers in Computational Neuroscience 12 (2018).

[81] E. Ullner, A. Politi, and A. Torcini, Quantitative and qualitative analysis of asynchronous neural activity, Physical Review Research 2, 023103 (2020).

[82] W. Bialek, M. DeWeese, F. Rieke, and D. Warland, Bits and brains: Information flow in the nervous system, Physica A: Statistical Mechanics and its Applications 200, 581 (1993).

