## Supplementary Information for "Switches in single-neuron excitability class can induce rich dynamical states in balanced networks"

### Information Theory

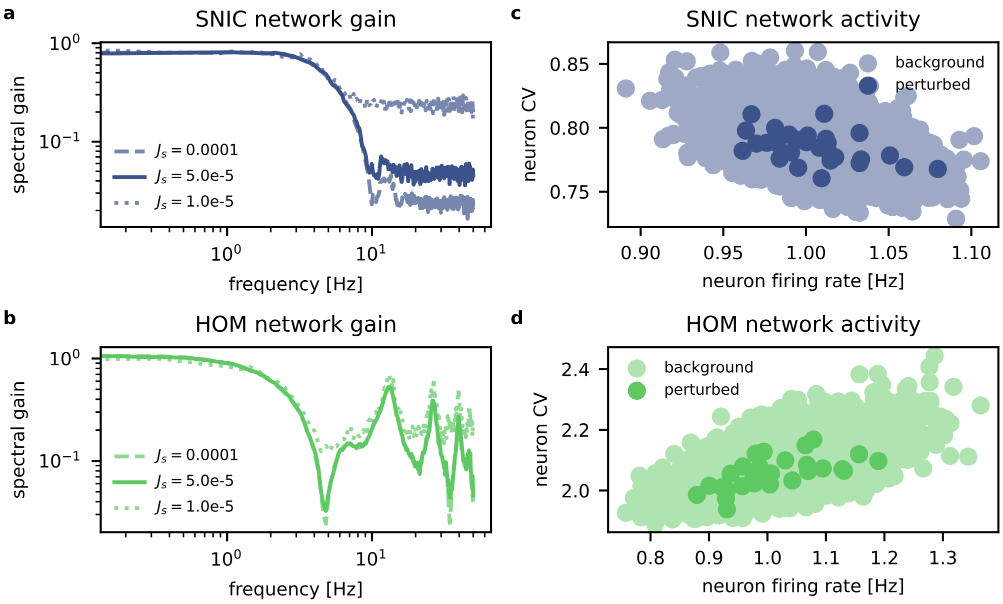


**Fig. S1 Confirmation of linearity for linear response. a) & b)** Spectral gain of the SNIC and HOM network for different signal strengths $J_s$. Input must be weak enough so that the gain remains constant but strong enough to reduce the noise level. For Fig. 2 (main manuscript), $J_\mathrm{s} = 5.0e - 5$ is used. **c) & d)** Firing rates and CVs of all SNIC (or HOM) neurons (light) and the stimulated neuron (dark) for 30 network realizations. Stimulated neuron's activity remains well within the distribution of the network. The simulated networks consist of $N = 500$ neurons, each with $K = 20$ inputs of strength $J_0 = 1$. The reset is $v_\mathrm{r} = -0.8$ for neurons in the SNIC network and $v_\mathrm{r} = 0.115$ for HOM. The network activity is tuned to $\nu = 1\,\mathrm{Hz}$. The simulation is run for $T = 2e7\,\mathrm{s}$.

### Chaos

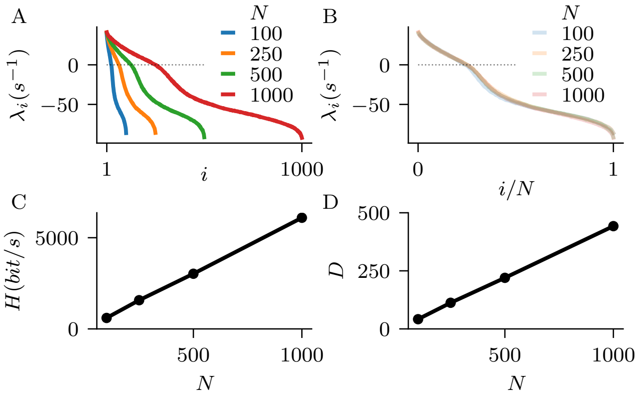


**Fig. S2 Extensive chaos in balanced networks in the HOM regime revealed by size-invariance of Lyapunov spectrum A)** Lyapunov spectra in the HOM regime for different network size $N$. **B)** The shape of Lyapunov spectrum, rescaled by network size $N$ is invariant of network size $N$, indicating extensive chaos. **C)** Dynamical entropy rate grows linearly with network size $N$. **D)** Similarly, the attractor dimension grows linear with $N$. Other parameters are $J_0 = -1$, $K = 20$, $\nu = 6\,\mathrm{Hz}$)

### Conductance-based models

While the present results based on the normal-form reduction of excitability classes are general with regards to other neuron models, we do reproduce the phenomenological effect using a reduced conductance-based Traub-Miles (TM) model [1, 2]. The model consists of a leak current, a sodium current and a potassium current

$$C\dot{v} = g_{\mathrm{L}}(E_{\mathrm{L}} - v) + \bar{g}_{\mathrm{Na}} m^3 h (E_{\mathrm{Na}} - v) + \bar{g}_{\mathrm{K}} n^4 (E_{\mathrm{K}} - v) + \sqrt{K} I_{\mathrm{ext}},$$

each with voltage-dependent gating kinetics

$$\begin{aligned}\dot{m} &= \alpha_m(v)(1-m) - \beta_m(v)m\\ \dot{h} &= \alpha_h(v)(1-h) - \beta_h(v)h\\ \dot{n} &= \alpha_n(v)(1-n) - \beta_n(v)n\end{aligned}$$

and Hodgkin-Huxley type activation curves.

$$\alpha_m(v) = \frac{0.32}{\mathrm{mV}} \cdot \frac{v + 54\,\mathrm{mV}}{1 - \exp\left(-\frac{v+54\,\mathrm{mV}}{4\,\mathrm{mV}}\right)} \cdot \frac{1}{\mathrm{ms}}$$

$$\beta_m(v) = \frac{0.28}{\mathrm{mV}} \cdot \frac{v + 27\,\mathrm{mV}}{\exp\left(\frac{v+27\,\mathrm{mV}}{5\,\mathrm{mV}}\right) - 1} \cdot \frac{1}{\mathrm{ms}}$$

$$\alpha_h(v) = 0.128 \cdot \exp\left(-\frac{v + 50\,\mathrm{mV}}{18\,\mathrm{mV}}\right) \cdot \frac{1}{\mathrm{ms}}$$

$$\beta_h(v) = \frac{4.0}{1 + \exp\left(-\frac{v+27\,\mathrm{mV}}{5\,\mathrm{mV}}\right)} \cdot \frac{1}{\mathrm{ms}}$$

$$\alpha_n(v) = \frac{0.032}{\mathrm{mV}} \cdot \frac{v + 52\,\mathrm{mV}}{1 - \exp\left(-\frac{v+52\,\mathrm{mV}}{5\,\mathrm{mV}}\right)} \cdot \frac{1}{\mathrm{ms}}$$

$$\beta_n(v) = 0.5 \cdot \exp\left(-\frac{v + 57\,\mathrm{mV}}{40\,\mathrm{mV}}\right) \cdot \frac{1}{\mathrm{ms}}$$

With the original model parameters

| | | |
|---|---|---|
| $C$ | $1\,\mu\mathrm{F}$ | Membrane capacitance |
| $E_\mathrm{L}$ | $-67\,\mathrm{mV}$ | Leak reversal potential |
| $E_\mathrm{Na}$ | $50\,\mathrm{mV}$ | Sodium reversal potential |
| $E_\mathrm{K}$ | $-100\,\mathrm{mV}$ | Potassium reversal potential |
| $g_\mathrm{L}$ | $0.1\,\mathrm{mS}$ | Leak conductance |
| $\bar{g}_\mathrm{Na}$ | $100\,\mathrm{mS}$ | Sodium conductance |
| $\bar{g}_\mathrm{K}$ | $80\,\mathrm{mS}$ | Potassium conductance |
| $V_\mathrm{Th}$ | $0\,\mathrm{mV}$ | Spike Threshold |

the neuron has a SNIC spike-onset. We induce the switch to HOM onset, by increasing the potassium reversal potential past the SNL point around $E_K = -63.5\,\mathrm{mV}$, again showing an increase in the spike time variability. The network parameters for this simulation are directly taken from Brunel 2000 [3], with $N_\mathrm{E} = 10000$ excitatory and $N_\mathrm{I} = 2500$ inhibitory neurons coupled randomly with a probability of $p = 0.1$, and a synaptic strength of $J_0 = 1.2\,\mathrm{mV}$. The network is tuned in the inhibition dominated regime by increasing inhibitory coupling $J_\mathrm{I} = -5J_\mathrm{E}$ beyond the balance according to subpopulation sizes. Excitatory synaptic strength $J_\mathrm{E} = J_0/\sqrt{pN_\mathrm{E}}$ is scaled with the average number of inputs. The connectivity consists of pulse-coupling, where the synaptic strength is added directly to the membrane voltage of the post-synaptic neuron. The connection probability leads to an average in-degree of $\bar{K} = 1250$ for each neuron, which is considerably higher than for the QIF networks used in the main paper. As examined in a recent study, networks of QIF neurons synchronize for high in-degrees [4], limiting our ability to examine different levels of sparseness for our main model. We therefore use this network of conductance-based neurons to show that neither the QIF neuron model, nor a low number of connections or the fixed in-degree connectivity are a requirement for the observed effect of burst-like activity. The dimensionality of this model unfortunately prohibits further analysis along the lines of the main paper. The firing rate of the networks is tuned to $\nu = 30\,\mathrm{Hz}$ by adjusting the external input current $I_\mathrm{ext}$, to account for activity-dependent effects.

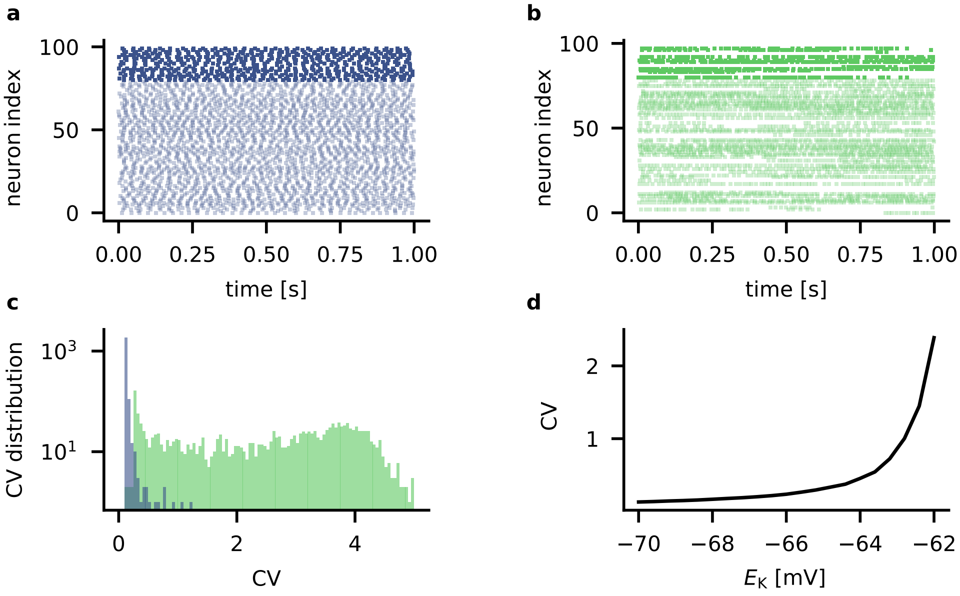


**Fig. S3 Network of conductance-based neurons.** A Brunel-type, inhibition-dominated balanced EI network with random connection probability and Traub-Miles neurons. **a)** Raster plot of asynchronous firing in the SNIC regime with $E_\mathrm{K} = -70\,\mathrm{mV}$, inhibitory (excitatory) neurons in dark (light). & **b)** Raster plot of the same network near the HOM regime for $E_\mathrm{K} = -62\,\mathrm{mV}$ (compare SNL switch at $E_\mathrm{K} = -63.5\,\mathrm{mV}$). **c)** Logarithmic histogram of the distribution of neuron CVs for both networks. **d)** The mean CV of the network for increasing potassium reversal potential $E_\mathrm{K}$. For model parameters, see text.

We further compare simulations of the described TM neurons with equivalent networks of Hodgkin-Huxley (HH) neurons which exhibit a Hopf spike-onset bifurcation [5, 6]. We add a bias input current of $I_\mathrm{bias} = 6.345\,\mu\mathrm{A}$ to the HH-Hopf neurons, to align their firing threshold with that of the TM-HOM neurons around $I_\mathrm{ext} = 0$. The network parameters for both networks are identical. They consist of $N_\mathrm{E} = 800$ excitatory and $N_\mathrm{I} = 200$ inhibitory neurons, each with a fixed in-degree of $K = 30$. The network is tuned in the inhibition dominated regime by increasing inhibitory coupling $J_\mathrm{I} = -5J_\mathrm{E}$ beyond the balance according to subpopulation sizes. Excitatory coupling $J_\mathrm{E} = J_0/\sqrt{K}$ is scaled by the in-degree and the networks are simulated for $T = 100\,\mathrm{s}$. The firing rate is not fixed in these simulations, as we explore a range of coupling strengths $J_0$ and external input currents $I_\mathrm{ext}$. For each combination of parameters, we calculate the firing rate and CV of the network as well as a measure of synchrony based on filtered spike-time cross-correlations [7]. Neurons with a Hopf spike-onset, being classified as class II excitability, also have a bistable input region. Beside other differences in their dynamics properties, we here demonstrate that a network of Hopf neurons is also capable of producing burst-like activity as we observe for HOM networks. The region of bistability for a Hopf neuron is much smaller, however, making it difficult for the neuron to be tuned to this regime. Further, the spiking irregularity as measured by the coefficient of variation is significantly lower and restricted to the direct surroundings of the spike onset, where these networks tend to synchronize (see Suppl. Fig.

S4). With careful tuning, however, a network of Hopf neurons does portray similar behavior to the HOM networks discussed in the main part of this study (Suppl. Fig. S5). Here, the network consists of $N_{\mathrm{E}} = 2400$ excitatory and $N_{\mathrm{I}} = 600$ inhibitory neurons, each with a fixed in-degree of $K = 200$. The model parameters for all Hopf neurons are as follows.

| | | |
|---|---|---|
| $C$ | $1\,\mu\mathrm{F}$ | Membrane capacitance |
| $E_{\mathrm{L}}$ | $10.613\,\mathrm{mV}$ | Leak reversal potential |
| $E_{\mathrm{Na}}$ | $115\,\mathrm{mV}$ | Sodium reversal potential |
| $E_{\mathrm{K}}$ | $-12\,\mathrm{mV}$ | Potassium reversal potential |
| $g_{\mathrm{L}}$ | $0.3\,\mathrm{mS}$ | Leak conductance |
| $\bar{g}_{\mathrm{Na}}$ | $120\,\mathrm{mS}$ | Sodium conductance |
| $\bar{g}_{\mathrm{K}}$ | $36\,\mathrm{mS}$ | Potassium conductance |

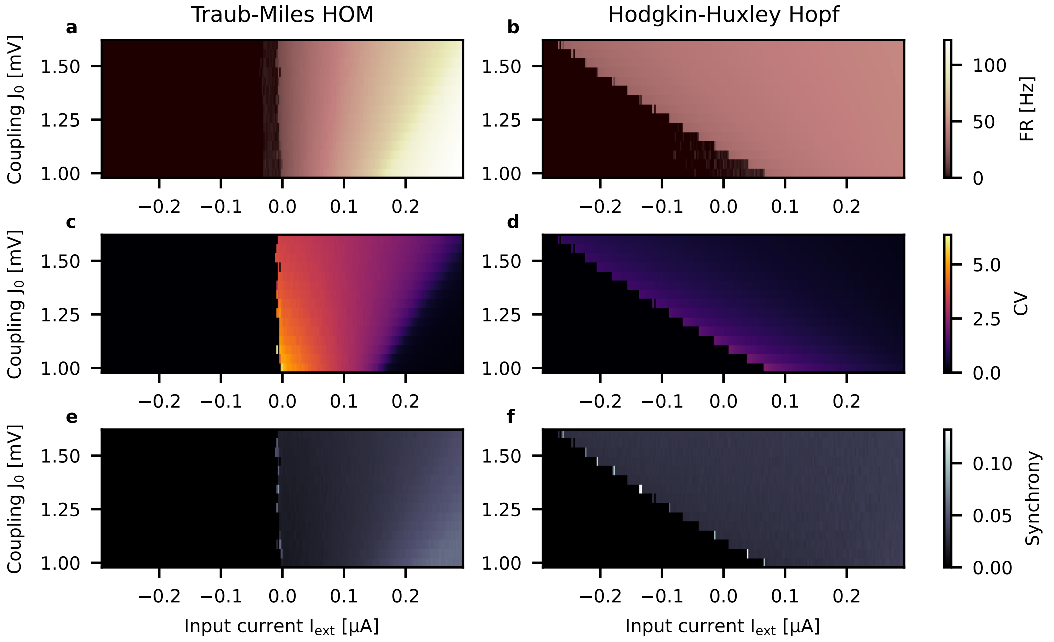


**Fig. S4 Comparison of HOM and Hopf network regimes.** Network simulations of conductance-based neurons for varying coupling strengths and input currents. **a)** & **b)** Colormap of the mean firing rate of the network for different coupling strengths $J_0$ and input currents $I_{\mathrm{ext}}$. **c)** & **d)** Mean coefficient of variation of each respective network. **e)** & **f)** Synchrony measure of each respective network. For simulation parameters, see text.

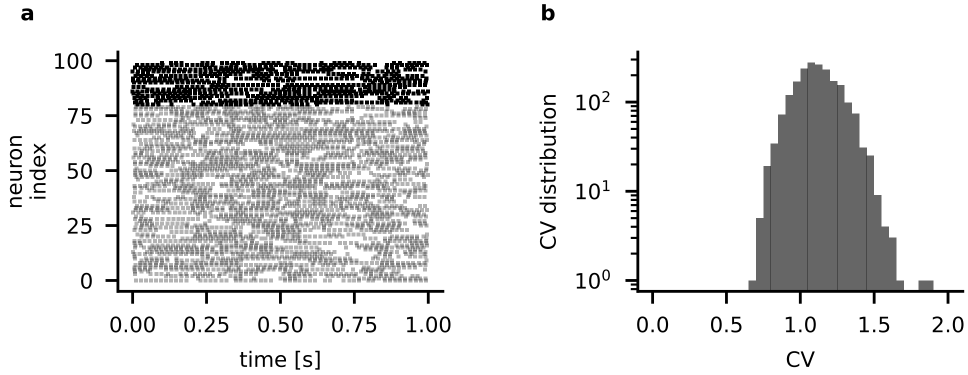


**Fig. S5 EI Network of Hodgkin-Huxley neurons with Hopf onset a)** Spike raster plot of the inhibitory (dark) and excitatory (light) population demonstrate burst-like activity in the network. **b)** The distribution of CVs in the network shows super-Poissonian irregularity.

#### *Influence of indegree on collective dynamics*

The degree of interconnectedness, meaning how sparse or dense a network is, has a fundamental influence on its dynamics. The model used in this study follows the sparse balanced networks analyzed by Brunel [3] in general, but more specifically the reduced version of the model used by Monteforte and Wolf [8]. As described in a more recent study, such networks of QIF neurons synchronize for high in-degrees [4], limiting our ability to examine different levels of sparseness for our main model and resulting in the low indegree used in the analysis. Notably, the synchronization with higher indegrees does not occur in networks of HOM neurons (Suppl. Fig. S6a-f), demonstrating the capability of neuronal HOM dynamics in decorrelating activity. Comparing the irregularity of the network states is not sensible once the SNIC network synchronizes. The CV of the HOM networks, while decreasing with higher indegrees (Suppl. Fig. S6h), remains super-Poissonian beyond the synchronization threshold $K \approx 130$ of the SNIC networks (Suppl. Fig. S6g).

For these simulations, the reset is set to $v_\mathrm{r} = -0.8$ for neurons in the SNIC network and $v_\mathrm{r} = 0.115$ for HOM. The network activity is tuned to $\nu = 1\,\mathrm{Hz}$. The simulated networks are modeled with increasing indegrees $K$, while the sparseness is kept constant by increasing the number of neurons as $N = 1000K$. Input strength is $J_0 = 1$.

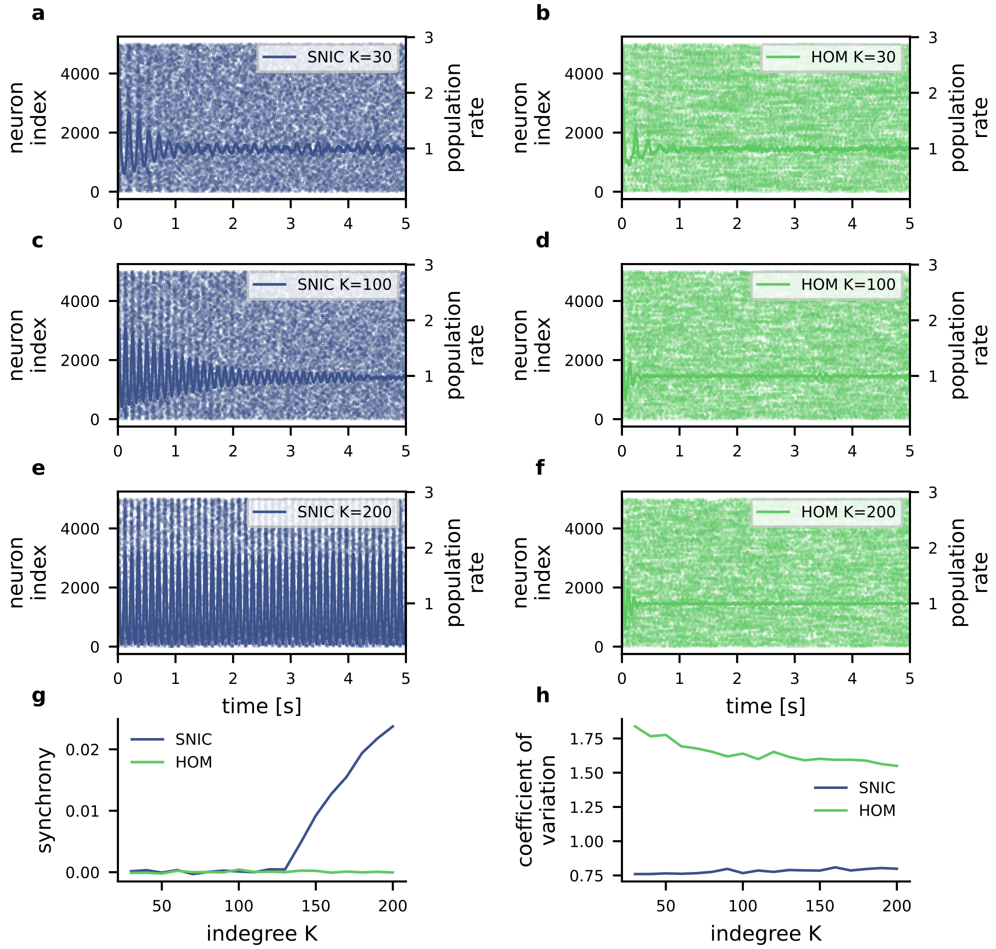


**Fig. S6 Emergence of network oscillations with increased indegree** For low indegrees, the SNIC network **a)** and HOM network desynchronize transiently from the start of the simulations. With increasing indegree, the oscillations decay slower in the SNIC networks **c)**, while the HOM networks quickly desychronize **d)**. Above a threshold of connectivity, the SNIC network displays stable oscillations **e)**, while the HOM network remains asynchronous **f)**. Network synchrony shows the transition point for SNIC networks at $K \approx 130$ **g)**. The CV remains largely stable through this synchronization in the SNIC networks **h)**, while it decreases with higher connectivity in the HOM case. For simulation parameters, see text.

---

[1] R. D. Traub, R. K. Wong, R. Miles, and H. Michelson, A model of a CA3 hippocampal pyramidal neuron incorporating voltage-clamp data on intrinsic conductances, Journal of Neurophysiology

**66**, 635 (1991).
[2] C. Börgers, An Introduction to Modeling Neuronal Dynamics, Texts in Applied Mathematics, Vol. 66 (Springer International Publishing, Cham, 2017).
[3] N. Brunel, Dynamics of Sparsely Connected Networks of Excitatory and Inhibitory Spiking Neurons, Journal of Computational Neuroscience **8**, 183 (2000).
[4] M. Di Volo, M. Segneri, D. S. Goldobin, A. Politi, and A. Torcini, Coherent oscillations in balanced neural networks driven by endogenous fluctuations, Chaos: An Interdisciplinary Journal of Nonlinear Science **32** (2022).
[5] A. L. Hodgkin and A. F. Huxley, A quantitative description of membrane current and its application to conduction and excitation in nerve, The Journal of physiology **117**, 500 (1952).
[6] E. M. Izhikevich, Dynamical Systems in Neuroscience (MIT press, 2007).
[7] S. Schreiber, J.-M. Fellous, D. Whitmer, P. Tiesinga, and T. J. Sejnowski, A new correlation-based measure of spike timing reliability, Neurocomputing **52**, 925 (2003).
[8] M. Monteforte and F. Wolf, Dynamical Entropy Production in Spiking Neuron Networks in the Balanced State Supplementary Material, Physical Review Letters **105**, 268104 (2010).
